# Digital polymerase chain reaction in an array of microfluidic printed droplets

**DOI:** 10.1101/860411

**Authors:** Yongfan Men, Jiannan Li, Tingting Ao, Zhihao Li, Bizhu Wu, Wen Li, Yi Ding, Kuo-Hao Tseng, Wen Tan, Baoqing Li, Yan Chen, Tingrui Pan

## Abstract

Digital polymerase chain reaction (PCR) is a fast-developed technology, which makes it possible to provide absolute quantitative results. However, this technology has not been widely used in research field or clinical diagnostics. Although digital PCR has been born for two decades, the products on this subject still suffer from either high cost or cumbersome user experience, hence very few labs have the willingness or budget to routinely use such product; On the other hand, the unique sensitivity of dPCR over traditional qPCR shows great potential applications. Here, a cost-effective digital PCR method based on a microfluidic printing system was introduced, trying to overcome those shortcomings. The microfluidic droplet printing technology was utilized in this study to directly generate droplet array containing PCR reaction solution onto the simple glass substrate for the subsequent PCR and imaging, which could be done with any regular flat-panel PCR machine and microscope. The method introduces a new perspective in droplet-based digital PCR in that the droplets generated with this method aligns well in an array without touch with each other, therefore the regular glass and oil could be used without any special surfactant. With simple analysis, the data generated with this method showed reliable quality, which followed the Poisson distribution trend. Compared with other expensive digital PCR methods, this system is more affordable and simpler to integrate, especially for those biological or medical labs which are in need for the digital PCR options but short in budget. Therefore, this method is believed to have the great potential in the future market application.

## 1. Introduction

Digital polymerase chain reaction (PCR) is an absolute quantitative technology of nucleic acid molecules based on Poisson distribution principle. This technology has already shown great application potential in rare mutation detection and absolute nucleic acid quantification^1–4^. Therefore, digital PCR has been applied in various fields such as biotechnology, cell biology, genetic engineering, forensic science, medical science, and drug discovery research^5–13^

The dispersion of nucleic acid sample is the first and most important step in digital PCR. Ideally, the DNA templates should be completely dispersed in separate reaction units (such as micro-pores or droplets)^14–15^. These tiny reaction units do not interfere with each other, therefore avoid the cross-contamination during the amplification. After the PCR thermal cycling, the fluorescence signals in each microsystem are detected. If there is a positive signal, it proves that there is a template molecule in the reaction unit originally^16–19^. So far, there are two main types of sample dispersion methods, one is microfluidic chip-based format^20–26^ and the other is the droplet-based format^27–37^. For example, Thermo Quant Studio™ 3D digital PCR system uses a silicon chip containing through-hole array to manually disperse the liquid, while Bio-Rad QX200™ digital PCR system is an efficient droplet-based platform method that generates droplets as the micro-reactors of digital PCR. Over the past two decades, the droplet based digital PCR, or ddPCR, showed greater potential than chip-based dPCR, proven by both the academic publications^38^ and industrial perspective^39^, and this trend is still on-going and will be more obvious in the near future. However, even the most successful ddPCR system still require additional instruments or operations for the following amplification step and analysis, which are very tedious and cumbersome, and require additional training and practice. Therefore, from the market point of view, the ‘perfect’ digital PCR system on the market is still yet to come, and optimization of the workflow of digital PCR with new technologies is worthwhile and has long been expected.

Here we report a novel digital PCR approach based on a microfluidic adaptive printing (MAP) system. The MAP system can realize accurate, automatic and adaptive droplet’s printing functionality driven by pneumatic force. The droplet array generated by this system is well aligned, with the droplets separated from each other, therefore normal oil could be used with no need for surfactant. The droplet array was generated on an extremely simple substrate, with which the PCR thermal cycling and the imaging could be performed directly, by flat-panel PCR machines and microscopes that are routinely available in bio- and medical-related labs. Not only the droplet printing chip and PCR substrate are simple in design and fabrication and low in cost, the whole operational workflow is also very straightforward, which needs almost no special training to grasp the gist of operation. Additionally, the sizes and coordinates of the droplets can be tuned accurately by simply controlling the printing parameters. Any lab that is interested in possessing the digital PCR ability could potentially use this method with affordable cost to generate useful digital PCR data that could outperform qPCR results.

## 2. Material and methods

### 2.1. The microfluidic adaptive printing (MAP) system

The MAP system can realize accurate, automatic and adaptive droplet’s printing functionality. Controlled by a home-developed software, this system can realize intermittent or continuous droplet printing. Both of the array pattern and droplet size could be customized by several parameters (including printing frequency, pulse width, air pressure, droplet interval, and the arrayed layout). Our system has ten independently controlled air valves so that it could drive ten individual microfluidic print channels.

The MAP-ddPCR system is mainly composed of several individual parts: a high-speed three-axis motorized stage module (HW-KIT2M, Thorlabs) for the droplet array forming and relative distance adjustment between the printing chip and the PCR substrate; an solenoid valve array (MH2D, Festo) with filtered, compressed air as the driving module; a holder structure to mount the chip and the tubing bundle to connect the valves with the chip air inlets, as the printing head.

### 2.2. Printing chip and digital PCR substrate

The printing chip was used to transform the PCR solution into droplets. The printing chip consists of two polydimethylsiloxane (PDMS) layers. The thick layer, consisting of a straight channel, a liquid chamber, an air inlet and a sample inlet, was fabricated by casting the PDMS onto a silicon wafer mold, cut into shape and punched with hole puncher; the thin layer was a PDMS film with a printing nozzle (~100 μm) that was laser cut (CO2, laser brand). The two layers of PDMS were bonded by plasma treatment with the alignment of the nozzle with the chamber. The wafer mold of the thick layer was fabricated by soft lithography with negative photoresist (SU-8, Micro Chem).

The PCR substrate was used as the substrate for receiving the droplets and form them into an array, and performing the PCR thermal cycling and the imaging. To prevent oil spill, the PCR substrate was fabricated by bonding laser-cut PDMS square wall frame (internal dimensions 45 mm × 70 mm × 2 mm) onto a normal glass slide (50 mm × 75 mm × 1 mm)

### 2.3. The optical detection apparatus

After the PCR amplification, the PCR substrate was transferred to an inverted fluorescence microscope (IX71, Olympus) for imaging. The imaging setup on the microscope consists of a 4X objective lens (Olympus), a CCD camera (DP72, Olympus), a 460/20 nm fluorescence excitation filter, and a FAM dye emission and acquisition filter at 532/30 nm. With a costumed controlling scripts based on the μManager software (University of California, San Francisco), the motorized stage moves with a fixed pace, along with the sequential images taken by the camera. Later, the photos were stitched into a single image and was processed. each field of view containing four droplets (1.6 mm x 1.2 mm, 41,890 pixels/droplet)

### 2.4. Image Processing and data analysis

A home-developed droplet analysis program, developed in MATLAB, was used to analyze data. Firstly, the background was subtracted by top-hat filter, and the contrast of the image was enhanced automatically to better differentiate the positive droplets from total droplets for the following steps. Next, the total droplets were identified and counted, and the grayscale and the coordinates of them were individually recorded. Then, a histogram of the droplets’ statistics was drawn to show the distribution of the droplet number versus grayscale. Usually there would be two major peaks in this histogram, as one represents the positive droplets at larger intensity, and the other peak indicates the ‘blank’ droplets at lower intensity. A threshold would then be determined to divide these two peaks, therefore divide all of the droplets into two groups. A counting of the positive droplets, based on the above process, could then be done, so that the copy number of the target DNA molecule could be determined.

As a middle-step of this image processing, the MATLAB program also displays the positive droplets with asterisk mark on the contrast-enhanced image, for the users to artificially check if the algorithm performs correctly.

### 2.5. Biological experiments

The PCR sample solution was printed onto an array of 1,000 droplets (40 x 25), with the droplet diameter around 250 μm. After the printing, 2-3 mL mineral oil (M8410, Sigma) was used to fully overlay the droplets to prevent the evaporation.

For digital PCR amplification, we prepared the standard sample solution with the final concentration of 2 to 128 aM using human genomic cDNA (Quantagene). PCR reaction mix consisted of 10 μL of TaqMan^®^ Fast Advanced Master Mix (2x, Thermo Fisher Scientific Inc.), 1 μL of TaqMan^®^ Gene Expression Assay (FAM dye) mix (20x) (GAPDH gene, Hs02758991_g1, Thermo Fisher Scientific Inc.), 1 μL of template solution, and 8 μL of RNase-free water (Takara). The experiment of each sample was performed more than three times on a newly prepared printing chip and PCR substrate. A commercial qPCR instrument (Q225, Quantagene) was used to perform similar PCR of the GAPDH gene of human genomic c-DNA template in order to validate the MAP-ddPCR results.

Digital PCR sample on the glass substrate was amplified in a flat-panel PCR machine (6335CH703993, Eppendorf AG 22331 Hamburg). 2-3 drops of mineral oil were placed onto the heating block to help with the heat transfer. PCR thermal cycling protocols were typically set as 40 two-step cycles (5 s at 95°C and 15 s at 60°C) with an initial denaturing step (60 s at 90°C, 60 s at 55°C, 60 s at 90°C, and 60 s at 55°C). The whole process could be completed within 45 minutes.

## 3. Results

### 3.1 The MAP system, operation of chip and substrate, and software interface

The MAP system (shown in Figure 1a) can generate custom droplet array with high quality. The chip was simply mounted in a 3-D printed holder, which is hanging above the PCR substrate on the x-y stage, and the distance between the nozzle of the printing chip and the PCR substrate (i.e. the fly distance for the droplets) is adjustable by the z-axis stage. The control electronics was embedded in the 3-D printed white box.

**Figure 1.**
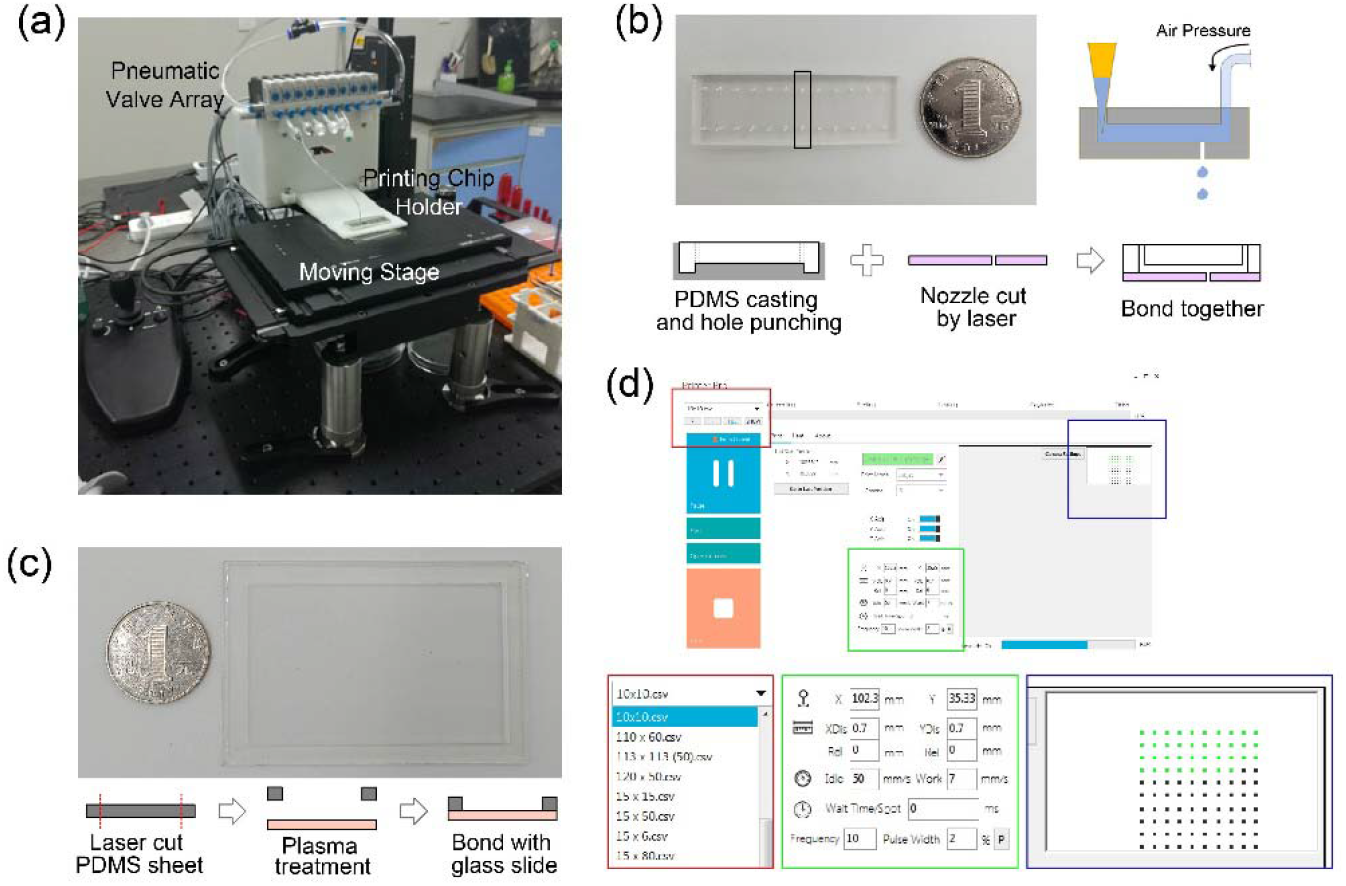
a) The MAP system, with the printing chip holder, 3-axis moving stage, and pneumatic valve array. b) The printing chip with 12 individual channels, with the inlet showing the printing principle, and the simple fabrication process. c) The PCR substrate, with the simple fabrication process. d) The software interface of the MAP system, with the ability to adjust the droplet array profile (red square), printing parameters (green square), and the progress monitoring (blue square).

As shown in Figure 1b, the printing chip (shown in the picture) has 12 simple-designed printing channels, with the nozzle beneath the straight microfluidic channel individually. The inlet on the right of the picture shows the operational protocol: firstly the regulated pneumatic pressure was guided into the 3-port, 2-way solenoid valve array that was connected to the chip inlet, and then the sample liquid was inserted into the entrance of the chip by pipette tip. Consequently, the printing system parameters were set through the computer printing software. The air pressure was regulated in a form of pulse train, resulting into a series of droplet formation from the nozzle. The fabrication process of the printing chip is shown below the picture, indicating that the top layer with the channel, the inlet, and the outlet was cast-molded, and the bottom layer with the printing nozzle was fabricated from a sheet by laser cutting; then, the two layers were bonded together to form a complete microfluidic chip.

The PCR substrate is shown in Figure 1c, with simple regular microscopic slide bonded with 2mm thick PDMS wall, to provide a substrate for the droplet array generation, PCR thermal cycling, and fluorescent imaging. After the printing, the mineral oil was applied to cover the droplet array to prevent the evaporation. The PDMS wall here plays an role of preserving the oil volume, making sure the droplet array is completely covered by the oil with no defect.

A software for the control of MAP system was also developed, and the software interface is shown in Figure 1d. Once the controlling software is opened, the computer will automatically find the correct serial port, connect, and feedback the connection status with every instrument, and move the 3-axis stages into the initial position set in the software. The users are allowed to set droplet array template (including the number of rows, number of columns, droplet number at each spot), as is shown in the red rectangle, printing mode (intermittent or continuous printing with non-stopping stage movement), and printing parameters (pressure, frequency, pulse width, air valve channel selection, and array step size), as is shown in the green rectangle. The printing process is also displayed throughout the printing period, as is shown in blue rectangle.

### 3.2 The characterization of the droplet array generated by the MAP system

As shown in Figure 2, the droplets generated by the MAP system was characterized for the size distribution and the linearity/ stability, and a scannable QR code was generated by dynamically controlling the printing the array template.

**Figure 2.**
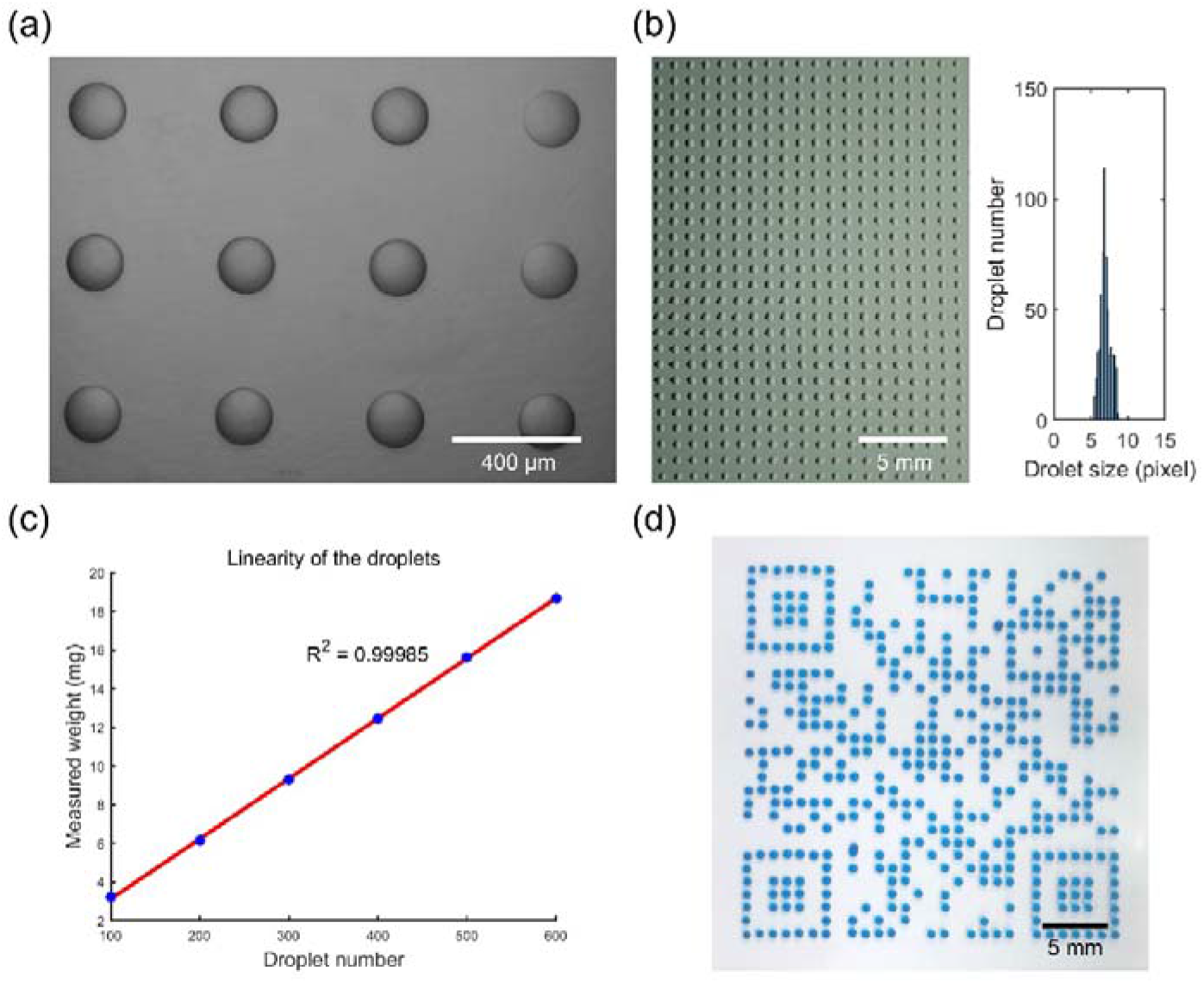
a) Microscopic image of printed droplet array. Scale bar is 400 μm. (b) Droplet size distribution analysis, showing a similar droplet size distribution and good monodispersity. Scale bar is 5 mm. c) The total weight measurement shows great linearity with the increase of the droplet number. d) A scannable QR code generated by the MAP system, indicating the flexibility of MAP system for other potential applications. Scale bar is 5 mm.

The droplets’ diameter is tunable from 50 μm to 500 μm, and in this paper, we used 250 μm droplets for the digital PCR experiments. A closer look, with a single view of the 4X objective lens, at the droplets printed on the PCR substrate and covered with oil shows great quality and distance (Figure 2a), that each droplet is perfectly dome shaped and the pacing among each other is in accordance with the design. The size distribution statistics (Figure 2b) indicates high level monodispersity and uniformity, with no defects among the array. The 0.9999 linearity of the droplet printing (Figure 2c) shows high stability as the droplets continuously being generated. The above three tests indicate that the droplet array generated by MAP system is stable and reliable over space and time. With such confidence, and with the flexibility of the MAP system, we generated a scannable QR code pattern (Figure 2d), which is a good example of the dynamic and precise control of the printing parameters and coordinates, and the flexibility of array template compatibility.

### 3.3 The workflow of the ddPCR experiments

As shown in Figure 3, with the MAP system, the dPCR sample solution, once prepared (Figure 3a) and loaded into the printing chip with a pipette tip, was directly printed on the glass substrates (Figure 3b). After the dispensing, the droplets were covered by mineral oil to prevent evaporation (Figure 3c), and then the PCR amplification was performed with optimized parameters (Figure 3d). Finally, the fluorescent signals were acquired by microscopy (Figure 3e) and proper processing and analysis was performed (Figure 3f).

**Figure 3.**
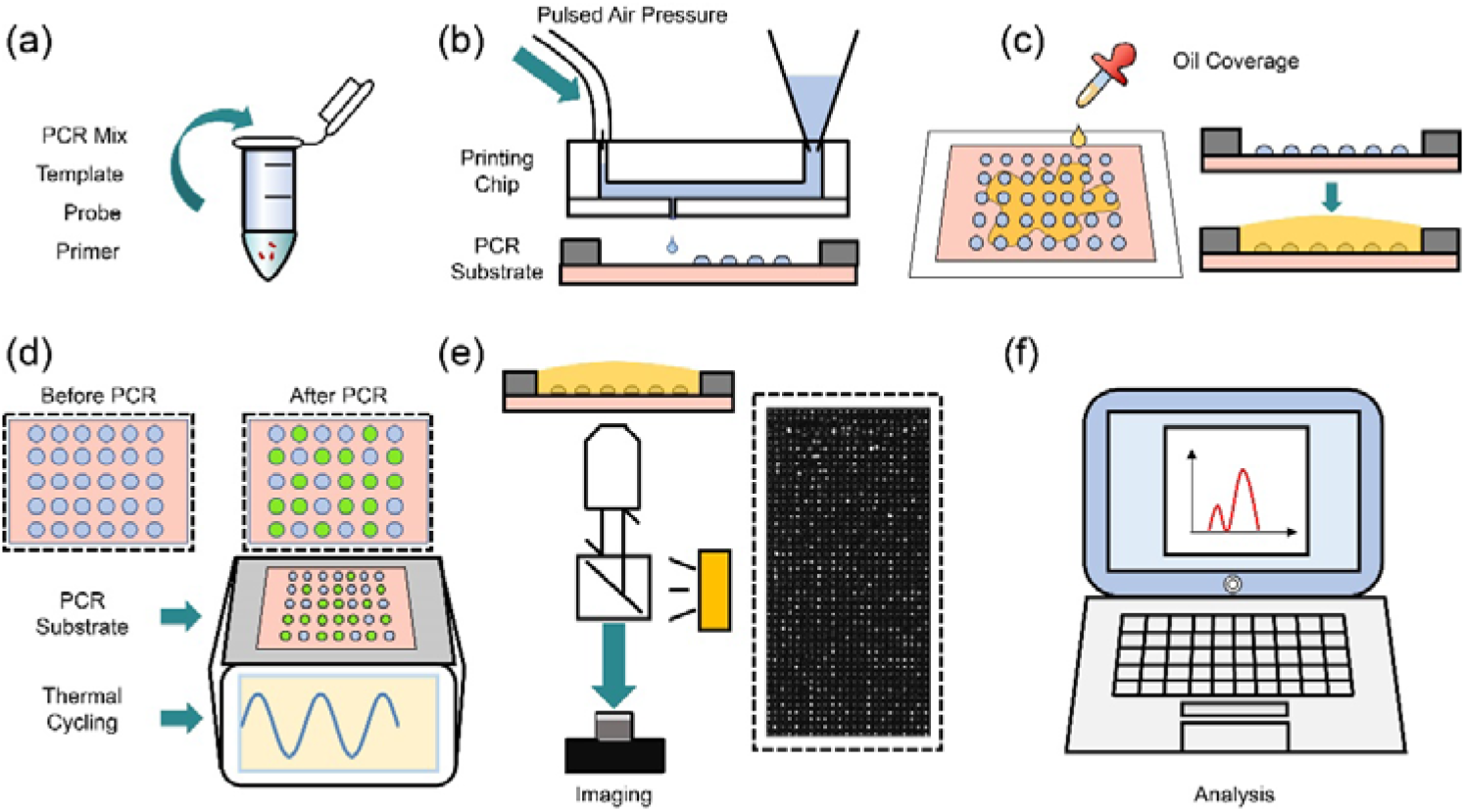
Digital PCR work flow based on MAP system. a) Sample preparation. The PCR mix, templates, probe, and primers were mixed in the centrifuge tube. b) Droplets printing. The sample solution is loaded into a pipette tip that is inserted in the inlet of the printing chip, and the other inlet of the chip is connected with the pneumatic valve. With the pulsed air pressure, droplets were ejected from the nozzle once at a time, and were collected on the PCR substrate. c) Oil loading. After the printing, mineral oil is applied above the droplet array to prevent evaporation. d) PCR thermal cycling. The PCR substrate was transferred to a flat-panel PCR machine for the PCR reaction. There is no extra pressure needed. e) Imaging. After the thermal cycling, the PCR substrate was transferred to an inverted microscope for imaging. The imaging process of the droplet array was performed in a photo-stitching manner with a custom-written script of the μManager software, for both bright-field and fluorescent channels. f) dPCR signal analysis. Different FOV images were stitched together into a complete fluorescent photograph for the following analysis, and the number of the positive droplets, as well as the negative droplets were analyzed, so that the absolute quantification of the target molecules was determined.

### 3.4 The results of the PCR amplification

Figure 4 shows the process needed to analyze the droplet image. First, the images were photo-stitched into a single raw image (Figure 4a). Second, the image was adjusted into higher contrast (Figure 4b and 4c, demonstrating only a portion of the whole image), in order to better determine the threshold for differentiate the positive droplets from the negative ones. After the contrast enhancement, the total intensity within each droplet was integrated, and a histogram was generated with the x-axis showing the intensity and the y-axis showing the statistical droplet number (Figure 4d). A good quality digital PCR data usually shows two individual peaks, with obvious gap (Figure 4d), so that the threshold (red dash line) could be determined with confidence and little error. Based on this threshold, positive droplets could be identified, marked, and counted (Figure 4e), as well as the negative droplets. After the Poisson correction, the absolute copy number of the template molecules could be calculated from the number of positive droplets, the signal percentage, and the droplet size, thus giving us an absolute quantification results.

**Figure 4.**
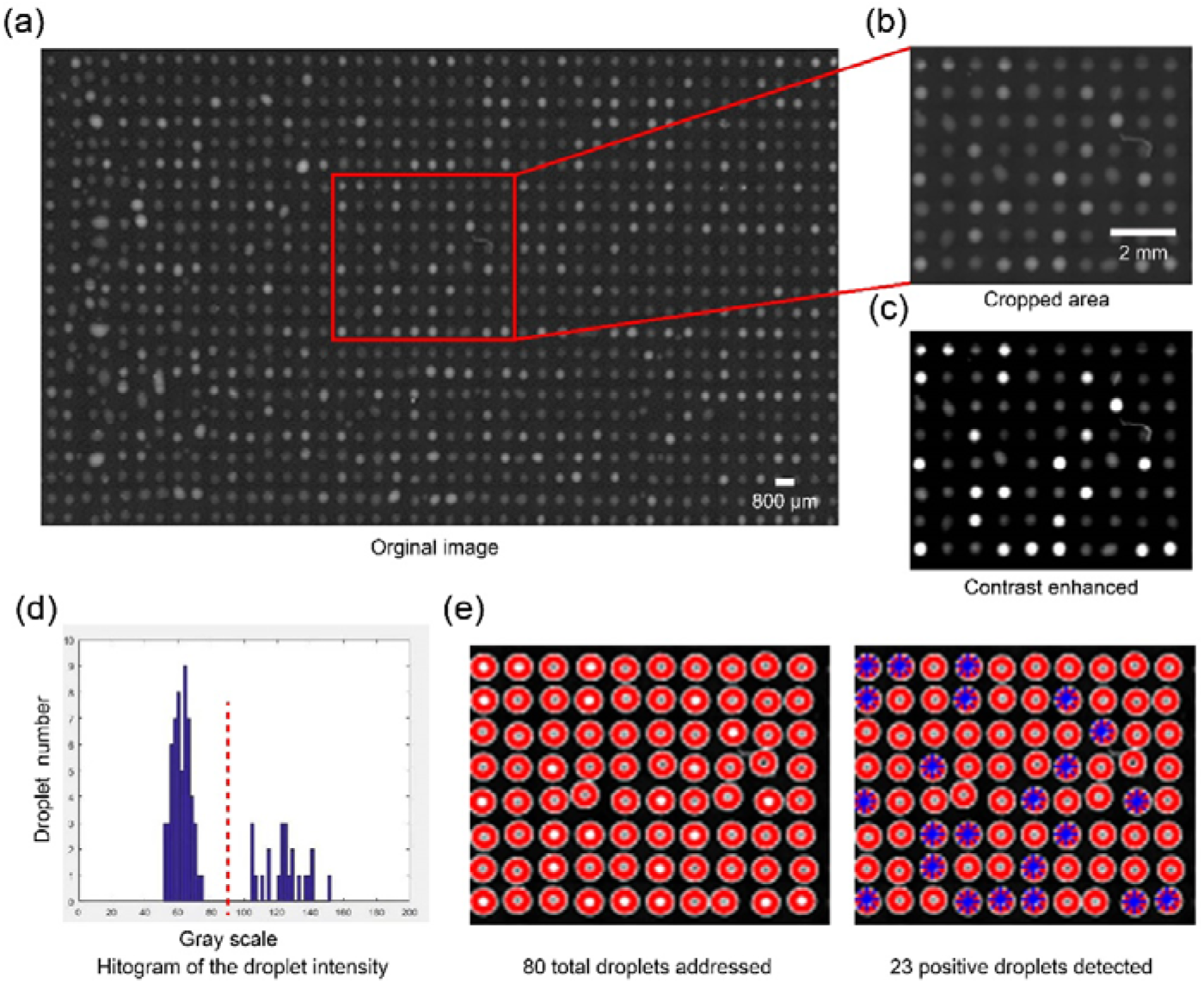
The digital PCR data analysis pipeline. a) Raw image (after photo stitching) containing 1000 droplets. Scale bar is 800 μm. b) A cropped region of the raw image as an illustrative example. Scale bar is 2mm. c) Positive droplets can be better distinguished from the negative ones after the contrast enhancement. d) A histogram of the droplet integrated intensity shows the threshold of the droplets could be easily determined, and the two groups could be clearly separated. e) The number and the coordinates of the total droplets, as well as the number of positive droplets were identified and marked.

### 3.5 The digital PCR results of the serial dilution

In order to evaluate the accuracy and reliability of the MAP system, a set of experiments of digital PCR were performed with seven different concentrations of standard human cDNA templates from 2 aM to 128 aM. Each sample was tested more than three times, and each experiment was performed on a newly prepared chip and substrate. As shown in Figure 5a, the average results of 7 groups of experiments were photographed. Each picture was processed as indicated in Figure 4, and the percentage of fluorescence signal was calculated. With the increase of template concentration, more positive droplets were observed under the fluorescence imaging. The bright droplets represented positive and they were distributed randomly indicating the digital PCR really happened. The result that the probability of the increase of positive droplets was consistent with the proportion of sample template concentration shown in Figure 5b proves that our system can easily get decent data that well-matched with predicted value in reference to Poison statics in different DNA samples, which shows our system’s accuracy in terms of DNA quantification.

**Figure 5.**
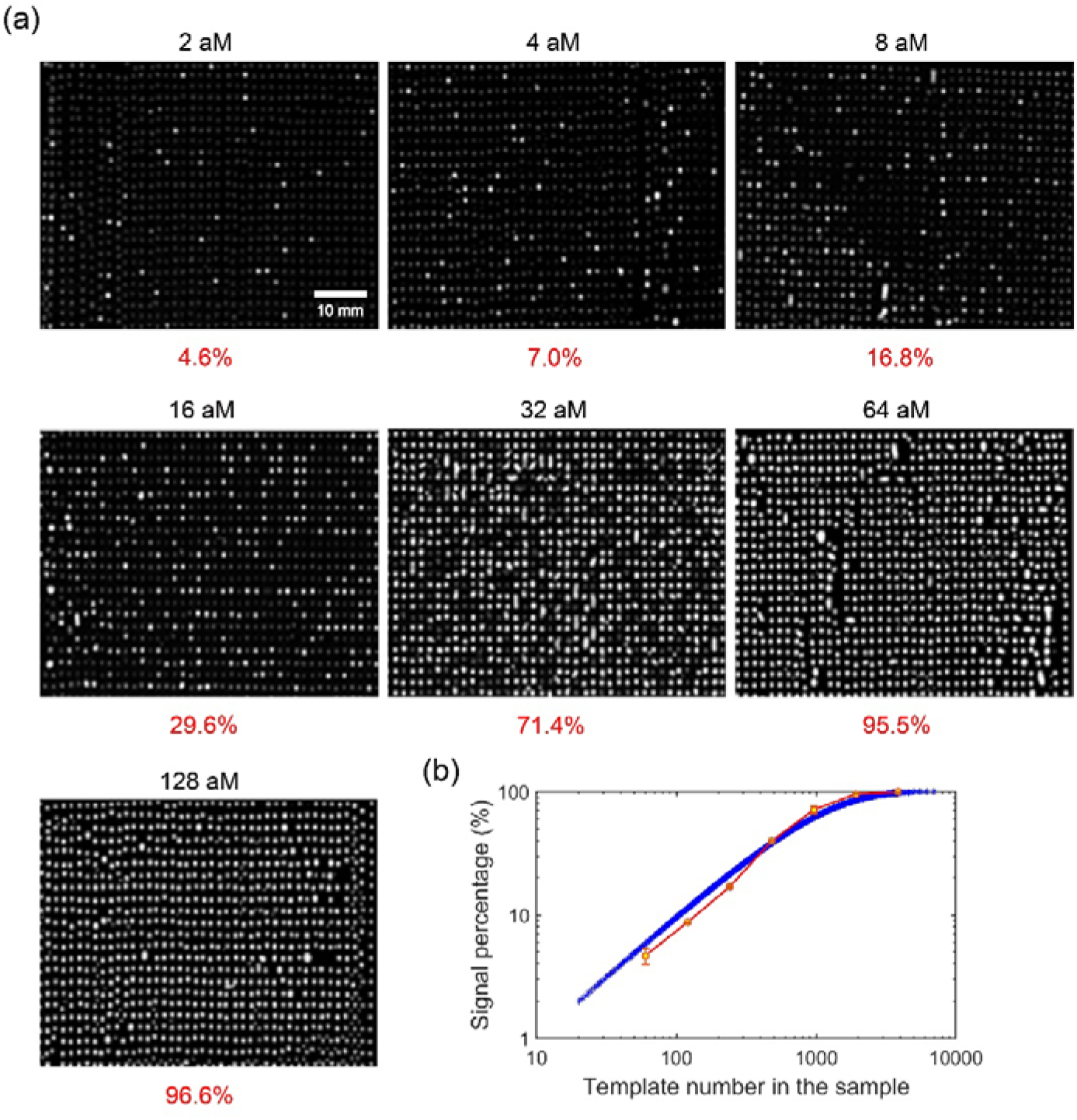
dPCR experimental results of serial dilution. (a) dPCR experiments of 7 different concentrations of templates, each concentration with three repetition (only one shown), was carried out. After the analysis, the percentage of the positive droplets of each experiment is shown. Scale bar is 10 mm. (b) The serial concentration dPCR results follows the Poisson distribution trend, and shows great linearity in the lower (<1000) copy numbers, therefore displays the ability of performing absolute quantification.

## 4. Discussion

The MAP system described in this paper provided the ability to generate customized droplet array, and the digital PCR data drawn from the droplets showed that the MAP system has a potential of a new digital PCR approach, adding to the current available methods in the field. To better illustrate our opinion, several sub titles are presented as follows:

i. **Advantages of MAP system.** The printing nature of the MAP system has several following advantages as opposed to other digital PCR methods: a) No needs to use special oil with surfactants, which may interfere with the PCR amplification and digital signals of the assay; b) The printing droplets form an array without any physical contact, therefore the possibility of cross contamination and droplet coalescence is erased; c) Coordinates of droplets are addressable, because once the droplet is printed on a spot, it won’t be moved again throughout the whole process. This property could trigger many possibilities, which will be talked about below. d) No enclosure needed around the droplets. Traditional digital PCR systems, no matter if it’s droplet based or chip based, we need to enclose the system during the thermocycling. Our method proves that in order to perform the PCR, no external pressure is needed, and just a casual mineral oil coverage is enough. This drastically reduces the requirement of the device. e) Low cost. The printing chip has relatively simple design and fabrication process that only requires single-layer photolithography, and there is no special treatment for the chip and the substrate, only regular cleaning would prepare the chip for reuse. If this technology were to be commercialized, both the printing chip and the PCR substrate could be fabricated by simple injection-molding, which would further lower the cost. The simple and low-cost property of the printing chip makes it disposable, to effectively avoid contamination.
ii. **Optimization potential:** the MAP system is eligible for the following improvements: a) The droplets in this paper were generated with 1 mm printing pace, but with the optimization of different parameters such as pressure, pulse width, distance between the printing chip and PCR substrate, we could realize even smaller droplets, and denser arrays. Therefore, even higher throughput, and larger dynamic range could be achieved. b) The printing chip is only made with PDMS; in the future, many other materials could be used to make the printing chip. For example, we have successfully used the plate sealing film as the bottom layer (data not shown). c) During the digital PCR experiment, only one channel was used, whereas the printing chip has 12 channels, therefore different types of solution could fill into different channels, and be printed based on demand, to optimize the reaction condition. For example, the PCR solution is usually mixed with PCR buffer, polymerase, MgCl_2_, primers, and probes. With multiple channels, we don’t need to pre-mix them anymore, and we could alter the combination ratio of the droplets among different rows and columns. Traditionally, we need to use multi-well plate to optimize the reaction concentrations, but now with MAP system, we could possibly regard the droplet array as the multi-well plate, and perform the reaction of thousands of droplets at once to screen out the optimized ingredient. d) The current MAP system is majorly for the droplet array generation, and the following digital PCR reaction and detection was carried out with the commercial-available devices, such as regular flat-panel PCR machine and inverted fluorescent microscope. All of the above stated modules together offer a low-cost, accessible, general solution that is affordable for any regular laboratory. Such solution provides characteristics such as cheap, easy to assemble, and universality across the regular lab. This is good for those bio-labs who have these regular machines available already, and could use the protocol in this paper to build a MAP system with relatively low-cost to realize dPCR function. e) However, from the user’s experience’ point of view, a push-button device would be better received. Following this thought, we have already integrated a peltier module to the MAP device for the temperature controlling (data not shown), therefore in the future, with the integration of the imaging module, the device would make it possible to perform the complete PCR reaction without moving the keep the PCR substrate. This might be the right direction for the commercialization of such technology. This down-stream integration would also present an opportunity to apply the technology to the biomedical detection applications.
iii. **Functionality expansion potential:** a) The above-stated properties of MAP can lead to many possible applications, such as adding content to the same droplet and single droplet tracking. We can even detect the fluorescent signal of each droplet while doing the thermal cycling. We could therefore do the qPCR on the droplet array with the integration of the image module and the heat block, which would become much better qPCR solution with a lot higher throughput and lower sample consumption than the current ones that are still based on 96-well plates. Especially combined with multi-volume technique, we could benefit from both qPCR and qPCR. b) The MAP system has customized user interface, which could realize several certain functions such as not only the droplet dispensing, but also the drop-to-demand and liquid adding on since the coordinates of the droplets are pre-determined. Therefore, MAP system could also realize accurately positioning and repetitive sample loading and buffer addition. Therefore, other than ddPCR, this MAP system also has many other possible applications. Because of this physical nature, our printing system is not only adaptive to dPCR, but also could be used for the following applications: single cell screening, single cell in-situ lysis and PCR amplification, and protein analysis^40–43^.
iv. **Disadvantages**. We have to admit that the MAP system also has some drawbacks/ shortcomings: a) The droplet size generated by MAP system is relatively large, which in turn consumes more PCR Mix if the droplet number is fixed. This is partially due to that the current printing parameters are optimized at this size for the best stability, and also due to that the PCR reaction results would be greatly suppressed when the droplets on the glass substrate are smaller than a certain size. We have a guess that the adsorbing of the enzyme on the glass surface is not negligible, but we still need more data to confirm. Ideally, an array of droplets could reach tens of thousands, whereas our array only has 1,000 droplets. By shrinking the size of the droplets, we may create much more droplets with the same volume of reaction mix. b) To prevent the droplets from further evaporating, the oil is applied after the droplets are printed, so the evaporation of the droplets before the oil is applied could cause the size of the droplets covered by oil quite different. We could design and fabricate the substrate, the could realize the oil be spread first, and then the droplet be printed. This could make the droplets in oil more uniform, therefore make the digital PCR results more reliable and trustable. c) Droplets are exposed to air for a short time as they are dispensed from the nozzle to the substrate. This leads to the problem of aerosol contamination, which has some limitations in biological safety and environmental protection. Printing process produced accompanied droplets, we adjusted the pressure and pulse width to avoid this situation. During the PCR thermal cycle, bubbles appeared and evaporated, which affected the fluorescence signal of surrounding droplets. d) There are several large droplets in some subset of Figure 5, such as in 64 aM and 128 aM. Those droplets are due to the fusion of droplets caused by the movement of the substrate from thermal cycler to the microscope for imaging, since there is no surfactant included in the oil, and the droplet is very flexible on top of the glass. This could be compensated by making patterned hydrophobic surface treatment onto the glass slide, or to integrate the whole facility together, so that there is no need to move the substrate.

## 5. Conclusion

In summary, an efficient and controllable microfluidic droplet printing platform is introduced. This platform can print droplet accurately and automatically. 1. **Precision** is reflected in that the system can print accurately according to the printing template (number of printing droplets, number of rows and number of columns) given by the user; 2. **High degree of automation** is reflected in that the system can automatically print according to the requirements of the experimenter after only adjusting the parameters; 3. **High adaptability** is reflected in the fact that the system can print different liquids from different channels. In addition, we also make **high-throughput** printing chips through PDMS to generate arrays of droplets of complicated QR code patterns. We have shown that this method can reduce sample consumption, control droplet size, and has high efficiency.

In the future, there will be other applications of our microfluidic droplet printing system. In addition to the digital PCR reaction, we can replace the sample droplets with single-cells. Single-cell printing has great prospects in biological applications. Linking the printing system to the microscope frees users from transferring from printing module to imaging module, which helps to monitor the printing process dynamically in real time. In addition to using PDMS, the thin layer of the printing chip in this study can be replaced by a more affordable single-sided adhesive sealing film.

## 7. Acknowledgements

The authors want to sincerely thank Yang Pan for the help of the discussion about the PCR experiments, and Bin Huang, for the help in the trial for the substrate surface modification, and Tiancong Zhao, who helped with the drawing of Figure 3. This research has been in part supported by National Natural Science Foundation of China (21904132), Program for Guangdong Introducing Innovative and Entrepreneurial Teams (2016ZT06D631), and the Fundamental Research Program of Shenzhen (JCYJ20180507182303606 and JCYJ20180302145552301)

## 8. Contribution list

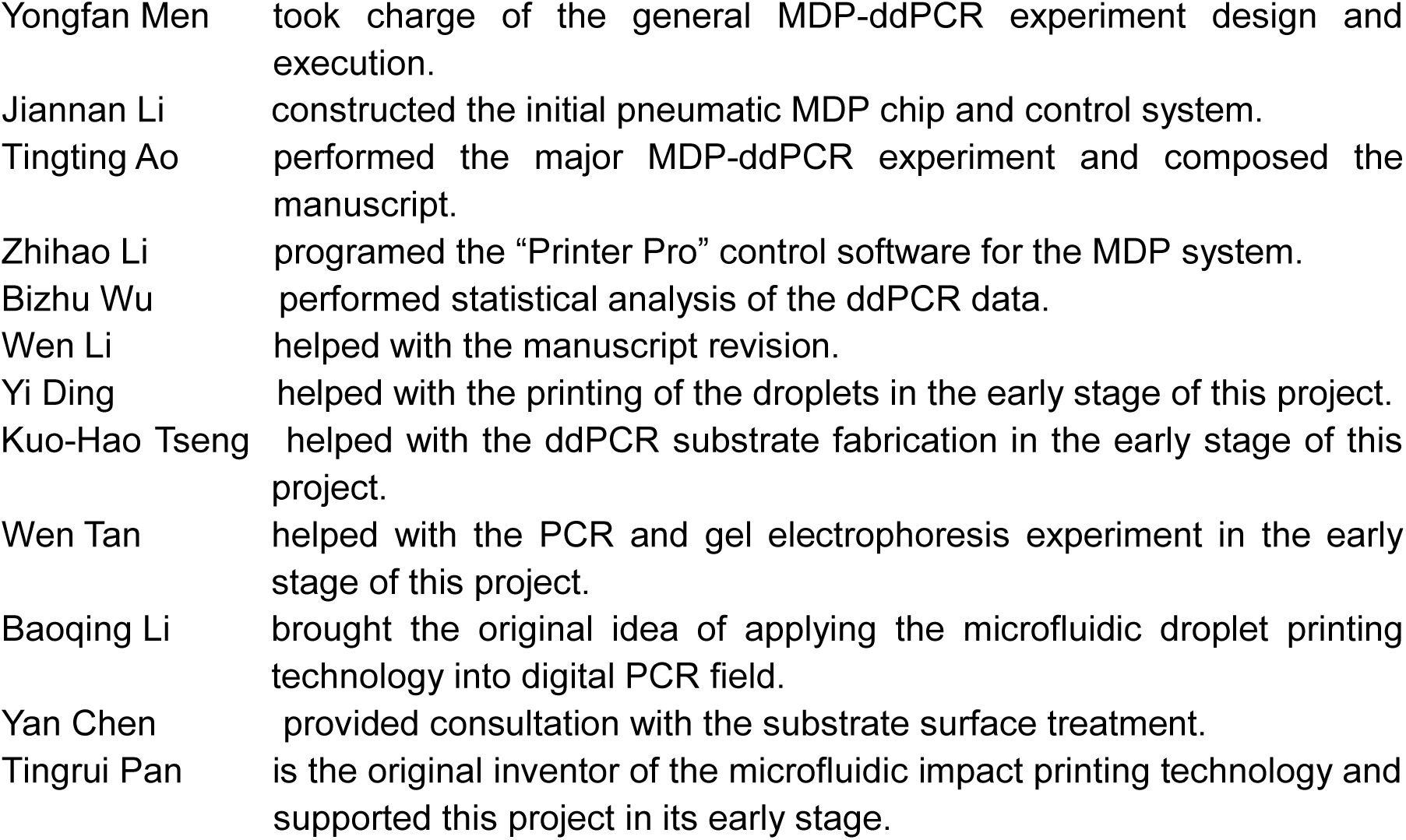

All authors have given approval to the final version of the manuscript.

